# Glial contribution to cyclodextrin-mediated reversal of cholesterol accumulation in murine NPC1-deficient neurons *in vivo*

**DOI:** 10.1101/2021.04.08.438990

**Authors:** Amélie Barthelemy, Valérie Demais, Izabela-Cristina Stancu, Eugeniu Vasile, Tom Houben, Michael Reber, Valentina Pallottini, Martine Perraut, Sophie Reibel, Frank W. Pfrieger

**Author notes:** These authors contributed equally (Amélie Barthelemy, Valérie Demais). Corresponding author or, INCI CNRS UPR 3212, 8 allée général Rouvillois, 67000 Strasbourg, France.

## Abstract

Niemann-Pick type C (NPC) disease is a rare and fatal lysosomal storage disorder presenting severe neurovisceral symptoms. Disease-causing mutations in genes encoding either *NPC1* or *NPC2* protein provoke accumulation of cholesterol and other lipids in specific structures of the endosomal-lysosomal system and degeneration of specific cells, notably neurons in the central nervous system (CNS). 2-hydroxypropyl-beta-cyclodextrin (CD) emerged as potential therapeutic approach based on animal studies and clinical data, but the mechanism of action on neurons has remained unclear. To address this topic *in vivo*, we took advantage of the retina as highly accessible part of the (CNS) and intravitreal injections as mode of drug administration. We find that CD enters the endosomal-lysosomal system of neurons and enables the release of lipid-laden lamellar inclusions, which are then removed from the extracellular space by specific types of glial cells. Thus, CD triggers a concerted action of neurons and glial cells to restore lipid homeostasis in the central nervous system.

## Introduction

Niemann-Pick type C (NPC) disease (OMIM #257220, OMIM #607625) is a rare, autosomal recessive and ultimately fatal lysosomal storage disorder with variable disease onset, multiple visceral and neurologic symptoms and diverging life spans (Bräuer et al., 2019; Gowrishankar et al., 2020; Hammond et al., 2019; Vanier, 2010; Wheeler and Sillence, 2020). The disease is caused by mutations in *NPC1* (Loftus et al., 1997) or *NPC2* (Naureckiene et al., 2000) that encode a membrane-resident and an intralumenal component of the endosomal-lysosomal system, respectively (Kwon et al., 2009; Li et al., 2017; Pfeffer, 2019; Qian et al., 2020; Winkler et al., 2019). Dysfunction of either protein causes accumulation of unesterified cholesterol and other molecules in late endosomes-lysosomes (Breiden and Sandhoff, 2020; Demais et al., 2016; Kobayashi et al., 1999; Liscum et al., 1989; Lloyd-Evans et al., 2008; Reid et al., 2004; Sokol et al., 1988; Zervas et al., 2001). The pathologic accumulation of cholesterol can be visualized by light microscopy following cyto- or histochemical staining with filipin (Pentchev et al., 1985), a bacteria-derived fluorescent polyene that binds unesterified cholesterol (Norman et al., 1972). Filipin staining of patient-derived fibroblasts has served as diagnostic test for the disease (Vanier and Latour, 2015). Electron microscopy revealed the presence of inclusions filled with lamellar membranes in neurons of NPC patients (Anzil et al., 1973; Harzer et al., 1978; Love et al., 1995; Palmer et al., 1985) and of animal models (Davidson et al., 2009; German et al., 2002; Lowenthal et al., 1990; Phillips et al., 2008; Praggastis et al., 2015; Tanaka et al., 1988).

At present, therapeutic options for NPC disease are limited, notably with respect to progressive neurologic symptoms (Geberhiwot et al., 2018; Toledano-Zaragoza and Ledesma, 2020). Several genetic and pharmacologic approaches are currently under study (Pallottini and Pfrieger, 2020). Cyclodextrins emerged as potential therapeutic approach based on a surprising finding: beneficial effects in NPC1-deficient mice originally attributed to allopregnanolone (Griffin et al., 2004) were in fact induced by the vehicle, 2-hydroxypropyl-beta-cyclodextrin (CD) (Davidson et al., 2009; Liu et al., 2008; Liu et al., 2009). Cyclodextrins are naturally occurring, water-soluble cyclic oligosaccharides with a cone-like shape that bind-and thereby solubilize-hydrophobic molecules (Coisne et al., 2016; Crini, 2014; Kurkov and Loftsson, 2013). Due to their size and the absence of a specific natural transport system, cyclodextrins hardly cross the intact blood-brain barrier (Banks et al., 2019; Camargo et al., 2001; Monnaert et al., 2004; Pontikis et al., 2013). Direct delivery of CD into the brain slowed neurologic disease progression in mouse and cat models (Pallottini and Pfrieger, 2020) of the disease (Aqul et al., 2011; Vite et al., 2015) and in human patients (Berry-Kravis et al., 2018; Farmer et al., 2019; Ory et al., 2017).

At present, it is unclear how CD accomplishes its beneficial effects in NPC disease *in vivo*. Previous studies investigated how CD affected NPC animal models and patients after chronic administration for weeks to months (Aqul et al., 2011; Berry-Kravis et al., 2018; Farmer et al., 2019; Fukaura et al., 2021; Ory et al., 2017; Palladino et al., 2015; Praggastis et al., 2015; Vite et al., 2015), but the immediate effects of CD on brain cells remain unknown despite growing interest in CD-based therapies for other neurodegenerative diseases (Coisne et al., 2016). Here, we addressed this fundamental question using the retina as accessible part of the CNS that is affected by NPC1 dysfunction in humans (Havla et al., 2020; Palmer et al., 1985) and in animal models (Claudepierre et al., 2010; Palladino et al., 2015; Phillips et al., 2008; Yan et al., 2014a). For drug administration, we chose intravitreal injections that enable direct delivery of molecules to the retina and subsequent monitoring of drug effects at defined time points (Del Amo et al., 2017). Vehicle injections in the contralateral eye establish the control condition in the same animal.

## Material and methods

### Animals

Experimental procedures involving animals and their care were performed in accordance with European and French regulations on the protection of animals used for scientific purposes (Directive 2010/63/EU and its transposition into the French regulation 2013-118, project authorised by the French Ministry of Research after ethical evaluation APAFIS#2016021212596886). All experiments were performed with 4-weeks-old Balb/c mice homozygous for the recessive NIH allele of *Npc1* or wildtype littermates (BALB/cNctr-Npc1^m1N^/J; Stock 003092; The Jackson Laboratory; Chronobiotron, UMS 3415, Strasbourg, France, ~90 generations of in-house matings). Mice were housed under SPF standard operating procedures (NPC colony positive for helicobacter sp, rodentibacter sp and mouse norovirus) in an individually ventilated caging system and at the following environmental conditions: 12/12h light/dark cycle (lights on at 7 am), temperature 21.5-23°C, 40-50% relative humidity. Autoclaved aspen wood chips (SAFE Lignocell Select) and nesting material (cotton squares and aspen chewing bars) were used as bedding environmental enrichment. Sterile-filtered or autoclaved tap water and irradiated standard rodent chow (SAFE A03-10, A04-10) were available ad libitum. Mice were genotyped as described (Buard and Pfrieger, 2014). Animals of either sex were used. At indicated ages and time points after injections, animals were deeply anethetized by intraperitoneal injections of 100 mg/kg ketamine and 20 mg/kg xylazine prior to dissection of eyeballs or retinae and euthanized by decapitation immediately after.

### Intravitreal injections

CD (100 mM; Sigma H5784) and vehicle (phosphate-buffered saline, PBS) were administered by intravitreal injections in 4-weeks-old animals. To this end, mice were anesthetized by isoflurane inhalation (3% for 15minutes) and injected using glass micropipettes and a micro-injector (Picospritzer III, Parker Hannifin, Pine Brooks, NJ, USA) in the right and left eye with drug and vehicle, respectively (volume 1 μl). One hour after injection, eyes were treated with gel to lubricate and rehydrate the cornea (Ocry-gel, TVM Lab) and mice were returned to their cage. For technical reasons, multiple intravitreal injections were not possible and observations were limited to a time period of 48 hours, before molecules are completely eliminated from the retina (Del Amo et al., 2017; Schmitt et al., 2019; Varadi et al., 2019).

### Production of CD-coupled nanoparticles

CD-gold (CD-AU) nanoparticles were prepared by fast reduction of the gold precursor chloroauric acid with sodium borohydride in the presence of CD. Using this method, cyclodextrin stabilizes nascent gold nanoparticles through hydrophobic-hydrophobic interactions without changing its binding properties (Cutrone et al., 2017; Liu et al., 2003). Briefly, a solution containing 10 mM CD in 0.25 mM HAuCl_4_ [50 mL; Gold(III) chloride hydrate, Sigma–Aldrich, Steinheim, Germany) was prepared in double distilled water (GFL bidistiller apparatus). Then, 100μL aliquots of freshly prepared 0.1M sodium borohydride (NaBH_4_) were added, at room temperature (RT; 20-24°C), until a stable orange colloid was noticed (1.2 mL of NaBH_4_). The reaction was allowed to proceed for 24h at RT. Resulting colloidal solutions were filtered using 0.45 μm Millipore filters, and stored at 4°C. CD-AU particles were deposited on a TEM copper grid covered with a thin amorphous carbon film with holes. High resolution transmission electron microscopy analyses of the films structure were performed using a TECNAI F30 G2 S-TWIN microscope operated at 300 kV and equipped with detectors for energy-dispersive X-ray (EDX) and electron energy loss spectrometry (EELS).

### Preparation, treatment and cytochemical staining of primary cultures

For *in vitro* experiments, RGCs were purified from one-week-old Wistar HAN rats (Chronobiotron, UMS 3415, Strasbourg, France) and cultured as described (Demais et al., 2016). Briefly, retinae were dissected and subjected to mechanic and enzymatic dissociation (Neural Tissue Dissociation Kit - Postnatal Neurons, Miltenyi Biotec). RGCs were purified using a subtraction (primary: rabbit anti-rat macrophage, Axell/Wak Chemie AI-A51240; secondary: goat anti-rabbit IgG, Jackson ImmunoResearch 111-005-003) and a selection step (primary: anti rat Thy1.1/CD90 clone T11D7e; secondary: goat anti-mouse IgM, Jackson ImmunoResearch 115-005-003) and plated on PDL-coated 96-well microplates (7000 cells per well; black/clear imaging plate, BD Falcon 353219). RGCs were cultured in serum-free medium (Neurobasal/Invitrogen) supplemented with MACS Neurobrew-21 (Miltenyi Biotec), brain-derived neurotrophic factor (25 ng/mL; PeproTech, London, UK), ciliary neurotrophic factor (10 ng/mL; PeproTech), forskolin (10 μM; Sigma), glutamine (2 mM; Invitrogen), N-acetylcysteine (60 μg/mL; Sigma), penicillin (100 units/mL; Invitrogen), sodium pyruvate (1 mM; Invitrogen), streptomycin (100 μg/mL; Invitrogen). RGCs were treated with glia-conditioned medium (GCM) after one day in culture. GCM was prepared as described (Nieweg et al., 2009). Briefly, mechanically dissociated cortices from P1-3 Wistar rats were cultured in poly-D-lysine (PDL)-coated (10 μg/mL; Sigma P7886) tissue culture plates (diameter 10 cm, BD Falcon Cat. 353003) in DMEM (#21969), heat-inactivated fetal calf serum (10%), penicillin (100 units/mL), streptomycin (100 μg/mL) and glutamine (2 mM) (all Gibco/Invitrogen). After two weeks, culture plates were washed with PBS and glial cells were cultured in Neurobasal supplemented with glutamine (2 mM), penicillin (100 units/mL), sodium pyruvate (1 mM), streptomycin (100 μg/mL) and NS21 (Chen et al., 2008). Two times a week, fresh medium was added and GCM was harvested after two weeks. GCM was spun down (5 min at 3000 g) to remove cellular debris. Two thirds of the RGC medium were replaced by GCM. To induce accumulation of unesterified cholesterol, RGCs were treated with 3-β-[2-(diethylamine)ethoxy]androst-5-en-17-one (U18666A; Interchim) after two days in culture at a concentration of 0.5 μg/mL for 48h. The drug induces cholesterol accumulation in cells (Karten et al., 2002; Liscum and Faust, 1989) by blocking NPC1 activity (Lu et al., 2015). Cultures were treated for 24h with indicated concentrations of CD [MW ~1400; 50 μM ~ 70 μg/mL ~ 0.007% (w/v)] and with CD-AU nanoparticles. For cytochemical detection of cholesterol, cultured RGCs were fixed by paraformaldehyde [PFA; 4% in PBS for 15 min at RT] and subjected to staining with filipin (40 μg/mL in PBS for 2 h at RT; from 250-fold ethanolic stock solution; F9765, Sigma). Fluorescence images of cultured cells were acquired using an inverted microscope (Axiovert 135TV; Zeiss) equipped with a metal halide lamp (10%; Lumen 200; Prior Scientific), an appropriate excitation/emission filter (XF02-2; Omega Optical Inc.), a 40x objective (oil, N.A. 1.3; Zeiss) and an air-cooled monochrome CCD camera (Sensicam, PCO Computer Optics) controlled by custom-written Labview routines (National Instruments). Micrographs were analysed semi-automatically by custom-written Labview routines (Demais et al., 2016).

### Histochemical and immunohistochemical staining

Eyeballs were rapidly removed from mice and immersion fixed using paraformaldehyde [PFA; 45 min in phosphate buffered saline (PBS) for 30 min] after an incision along the ora serata to remove the cornea. For whole-mount staining, retinae were dissected and processed in 96-well plates. For staining of retinal sections, eyeballs were embedded in agarose (5% in PBS), and cut at 50 μm thickness on a vibratome (Leica, VT1000S). For histochemical staining of unesterified cholesterol, retinal tissue (vibratome sections or whole-mounts) was incubated for 1h with filipin (2 μg/ml in PBS). For immunohistochemical stainings, retinal tissue was blocked (BSA 2%, goat serum 5% in PBS) and permeabilized (0.1% Triton with 0.1% BSA in PBS) for 45 min and incubated overnight at 4°C with primary antibodies (diluted in 1% BSA, 1% goat serum in PBS). Sections were washed and incubated for 2 hr at RT with secondary Alexa-labelled antibodies (1:500 in PBS). The following antibodies were used for immunohistochemical staining: anti-Tuj1 (1:1000; Covance; MMS-435P-250), anti-IBA1 (1:500; Wako; 019-19741), anti-GFAP (1:200; Sigma; G3893), anti-CD68 (1:100; Biorad; MCA1957), anti-MPO (1:500; Dako; A0398). For nuclear staining, retinal sections were incubated for 10 minutes with Sytox Green (0.25 μM; Invitrogen). Fluorescence images of retinal sections or whole-mounts were acquired on a confocal microscope (Leica SP5 II, Leica Microsystems) or an inverted microscope (Zeiss Observer 7) equipped with objectives (40x water, N.A. 1.2; 63x oil, N.A. 1.4), a module for optical sectioning by structured illumination (Zeiss ApoTome.2) and a digital camera (Hamamatsu ORCA-Flash 4.0). Micrographs were analysed semi-automatically using ImageJ (National Institutes of Health, Bethesda, MD) and custom-written Labview routines (Demais et al., 2016).

### Transmission electron microscopy

For ultrastructural analyses, retinae were fixed with glutaraldehyde (2.5% in 0.1 M cacodylate buffer at pH 7.4 for 2h at RT; Electron Microscopy Services-EMS) and rinsed with cacodylate buffer. Cholesterol was stained by filipin (50 μg/ml for 30 min, from a 100-fold dimethylformamide stock solution). Retinal sections were postfixed for 1h with osmium tetroxide [OsO_4_; 2%(w/v)] in 100 mM imidazole buffer to enhance lipid preservation before dehydration, embedding, and ultramicrotomy. To detect CD-AU particles, retinal sections were fixed with glutaraldehyde (2.5% in PBS at pH 7.4 for 1h at RT; Electron Microscopy Services-EMS), rinsed with PBS and then with double-distilled water. Gold particles were silver enhanced using the R-Gent SE-EM kit (Aurion). Sections were postfixed in OsO_4_ (0.5% in aqua bidestillata for 15 min), dehydrated in graded ethanol series and embedded in Embed 812 (EMS). Ultrathin sections were cut with an ultramicrotome (Leica), stained with uranyl acetate [1% (w/v) in 50% ethanol], and examined with a Hitachi H7500 transmission electron microscope equipped with a digital camera (Advanced Microscopy Techniques/Hamamatsu).

### 3β-hydroxy 3β-methylglutaryl Coenzyme A reductase (HMGCR) activity assay

The assay was performed by the radioisotopic method based on the production of [^14^C]MVA from 3-[^14^C]-HMG-CoA (specific activity 57.0 mCi/mmol., Amersham-Pharmacia, Little Chalfont, UK) as described (Segatto et al., 2014). Briefly, retinae were homogenized (1:5 w/v) in PBS containing (in mM) 100 sucrose, 50 KCl, 40 KH_2_PO_4_, 30 EDTA, pH 7.4 and incubated in the presence of co-factors (20 mM glucose-6-phosphate, 20 mM NADP sodium salt, 1 unit of glucose-6-phosphate dehydrogenase and 5 mM dithiothreitol). The assay was started by the addition of 3-[^14^C]-HMG-CoA (0.088 μCi/11.7 nmol). The newly synthesised [^14^C]-MVA was purified by ion exchange chromatography (AG1-X8 resin; BioRad Laboratories, Hercules, CA, USA) and its radioactivity was measured by a liquid scintillation counter (Perkin Elmer 2100TR). An internal standard (3-[^3^H]-MVA, specific activity 24.0 Ci/mmol., Amersham-Pharmacia, Little Chalfont, UK) was added to calculate recovery. HMGCR activity was expressed as pmol/min/mg protein.

### Tissue lysate preparation and immunoblotting

Total lysate of retinae was obtained by tissue homogenization in 1:5 w/v homogenization buffer (in mM 10 Tris-HCl, 1 CaCl_2_, 150 NaCl, 1 PMSF, pH 7.5). The samples were sonicated (VCX 130 PB, Sonics, Newtown,06470 CT) on ice for 30 sec, and centrifuged for 7 min at 13,000 rpm at 4 °C to yield total lysate, which was boiled for 5 min before SDS-PAGE and subsequent immunoblotting. Proteins were separated by SDS–PAGE and blotted to nitrocellulose membranes (Trans-blot Turbo, BioRad). Immunoblots were incubated with anti HMGCR primary antibody (Upstate, Lake Placid, NY; 1:500) overnight, followed by secondary peroxidase-conjugated antibodies produced in rabbit (1:10,000; Biorad). Immunoreactivity was detected by enhanced chemiluminescence (GE Healthcare, Little Chalfont, United Kingdom). As loading control, the immunoblots were reacted with an antibody against tubulin (1:10,000; Sigma Aldrich). Band intensities were quantified using ImageJ (National Institutes of Health, Bethesda, MD).

## Results

We studied the immediate effects of CD on retinal neurons *in vivo* using intravitreal drug injections and NPC1-deficient BALB/c mice at 4 weeks of age before animals showed neurologic symptoms.

### Pathologic cholesterol accumulation in retinal neurons of NPC1-deficient mice

First, we examined the distribution of unesterified cholesterol in retinae of 4-weeks-old mutant (NP) mice. Previous studies noted lipid accumulation in the retina of NPC mice at two months of age (Claudepierre et al., 2010; Palladino et al., 2015; Yan et al., 2014a), but the situation in younger animals has not been described. Cytochemical staining of retinal sections revealed the presence of filipin-positive puncta in all layers of four-weeks-old NP mice, and their absence from retinae of wildtype (WT) littermates. Notably, the density of filipin-positive puncta showed a high-to-low gradient from the inner to the outer retina with the ganglion cell and amacrine cell layers showing the strongest cholesterol accumulation (Fig. 1A). Previous studies reported the presence of lamellar inclusions in neurons of the ganglion cell layer (GCL) in NPC patients (Palmer et al., 1985) and NPC1-deficient mice (Claudepierre et al., 2010; Yan et al., 2014a). Moreover, we and others demonstrated that these structures represent the ultrastructural correlate of lipid accumulation (Demais et al., 2016; Kwiatkowska et al., 2014). Inspection of retinal sections by transmission electron microscopy (TEM) confirmed the presence of lamellar inclusions in retinal neurons from NP mice and their absence from retinae of WT littermates. Notably, we observed layer-specific differences: Neurons in the GCL had more and larger inclusions than those in the amacrine cell layer (ACL; Fig. 1C). In line with neuron-specific transcriptional changes due to NPC1 deficiency (Demais et al., 2016), the disturbed cholesterol flow within retinal cells induced an upregulation of cholesterol synthesis as indicated by enhanced activity and protein level of the rate-limiting enzyme, HMGCR (Fig. 1D). Together, these results revealed layer-specific accumulation of cholesterol in retinal neurons of presymptomatic mice and thus validated their use to explore the effects of CD.

**Fig. 1.**
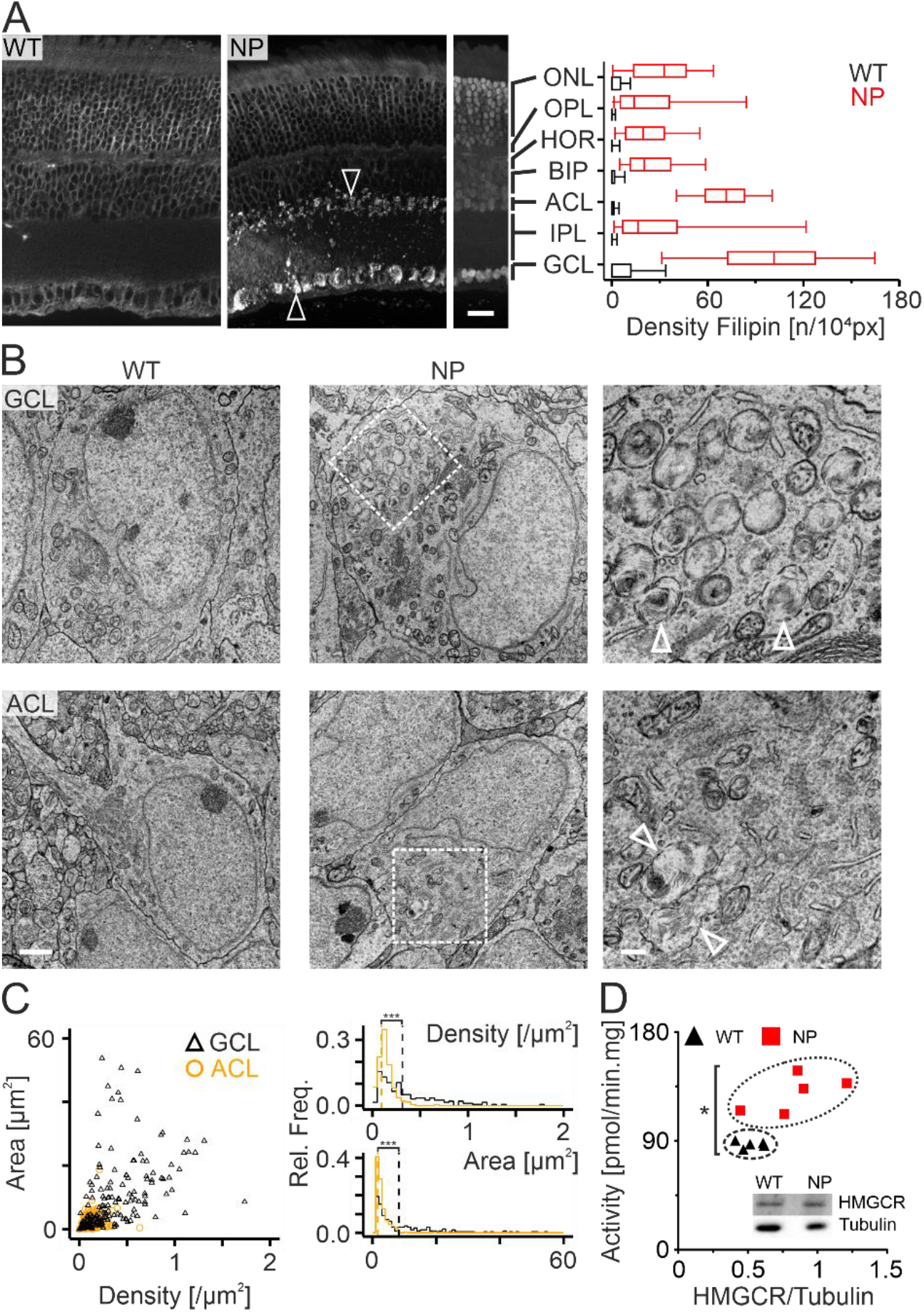
Pathologic accumulation of cholesterol in retinae of NPC1-deficient mice. A, Fluorescence micrographs (left) of retinal sections from 4-weeks-old wildtype (WT) and mutant mice (NP) subjected to staining with filipin and Sytox green. Arrowheads (left) indicate cells in the GCL and ACL with strong accumulation of unesterified cholesterol. Scale bar: 50 μm. Quantitative analysis (right) showing densities of filipin-positive puncta in indicated retinal layers of WT and NP mutant mice. Asterisks indicate statistically significant differences between WT and NP mice (p < 0.0001; ANOVA with Tukey post-hoc test; 37 - 680 ROIs from 4 WT or NP mice). ONL, outer nuclear layer; OPL, outer plexiform layer; HOR, horizontal cells; BIP, bipolar cells and Müller cells; ACL, amacrine cell layer; IPL, inner plexiform layer; GCL, ganglion cell layer. B, Transmission electron micrographs showing representative neuronal somata in the GCL and ACL from WT and NP mice. Note the presence of lamellar inclusions (indicated by arrowheads) in NPC1-deficient neurons. Scale bars: 2 μm (left, middle), 500 nm (right). C, Scatterplots (left) and relative frequency histograms (right) of the density and the area of inclusions present in neuronal somata of the GCL (black, NP: 5 animals / 200 cells) and of the ACL (orange, NP: 5 / 248) from NPC1-deficient retinae. Asterisks indicated statistically significant differences between the two layers (p < 0.001; Wilcoxon signed rank test). D, Scatterplots of enzyme activity and normalized protein levels of HMGCR in retinae from WT and NP mice indicating an increase of cholesterol synthesis due to NPC1-deficiency. Insert, representative immunoblot showing levels of HMGCR and tubulin. Asterisk indicates statistically significant increase in enzyme activity due to NPC1 deficiency (p < 0.05; Wilcoxon signed rank test; n = 5 mice).

### Reaction of retinal glia to NPC1-deficiency

Previous studies showed activation of glial cells in the CNS of NPC patients and animal models with distinct cell type- and region-specific onsets (Baudry et al., 2003; Cologna et al., 2014; Cougnoux et al., 2018; Cougnoux et al., 2020; Gabande-Rodriguez et al., 2019; German et al., 2002; Kavetsky et al., 2019; Lopez et al., 2011; Luan et al., 2008; Maue et al., 2012; Maulik et al., 2012; Park et al., 2019; Pressey et al., 2012; Repa et al., 2007; Santiago-Mujica et al., 2019; Seo et al., 2014; Stein et al., 2012; Tanaka et al., 1988; Walterfang et al., 2020; Yan et al., 2014a; Yan et al., 2014b). We asked how retinal glial cells react to the absence of NPC1 in 4-weeks-old mice. Immunohistochemical staining of retinal whole-mounts revealed increased densities and areas of IBA1-positive microglial cells in retinae of NP mice compared to those from WT littermates. However, no changes were observed for GFAP-positive cells at this age (Fig. 2A, B). These findings are in line with observations that microglial activation precedes the reaction of GFAP-positive cells in brains of NPC mutant mice (Baudry et al., 2003; German et al., 2002; Maulik et al., 2012; Park et al., 2019). Next, we examined the ultrastructural changes in glial cells of the GCL and ACL, where neurons showed strong cholesterol accumulation. The fractional area covered by Müller cells and glial cells with clear cytoplasm (Fig. 2C) was not affected by NPC1 deficiency in four-weeks-old animals, whereas the small fractional area covered by cells with dark cytoplasm was further decreased in the GCL (Fig. 2D). Based on established ultrastructural criteria (Bisht et al., 2016; Kavetsky et al., 2019; Peters et al., 1991; Ramírez et al., 1996) cells with clear and dark cytoplasm correspond to astrocytes and microglial cells, respectively. Lamellar inclusions were absent from retinal Müller cells in NP mice (total of 2410 μm^2^ and 1486 μm^2^ for n= 6 WT and n = 6 NP mice) in line with a previous study (Palmer et al., 1985). They were also not detectable in glial cells with clear and dark cytoplasm, which covered relatively small areas of the GCL and ACL regardless of the genotype. Together, these results revealed minor changes in retinal glial cells of NPC1-deficient presymptomatic mice.

**Fig. 2.**
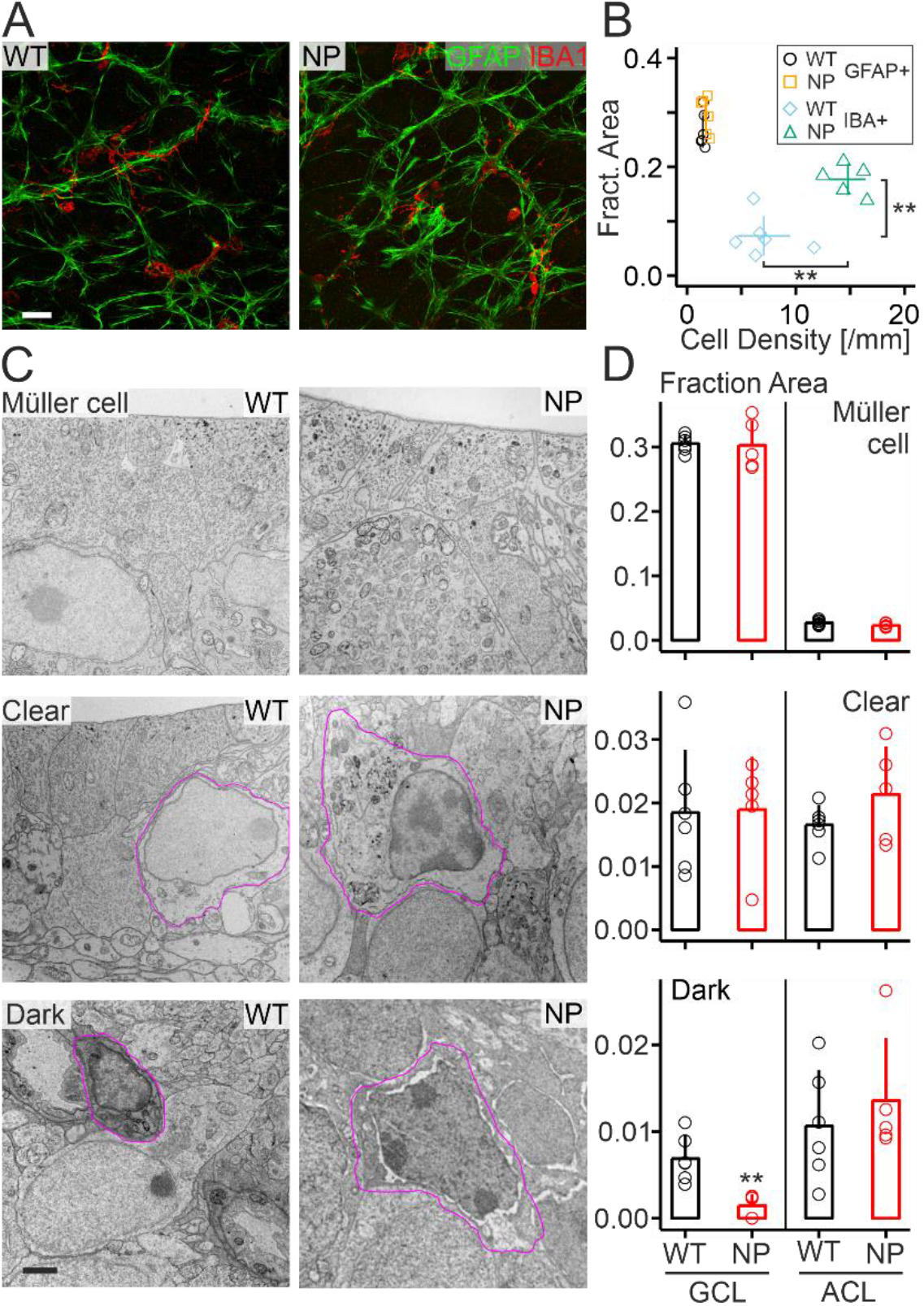
Glial reaction to NPC1 deficiency in retinae of 4-weeks-old mice. A, False-color fluorescence micrographs showing the distribution of indicated markers in retinal whole-mounts from WT and NP mice. Scale bar: 25 μm. B, Scatterplots of the mean fractional area and of the mean density of glial cells positive for the indicated markers in individual mice with the indicated genotype. Asterisks indicate statistically significant changes (p < 0.01; Wilcoxon signed rank test). C, Electron micrographs of Müller cells and glial cells with clear and dark cytoplasm in retinae of WT and NP mice. Scale bar: 2 μm. D, Mean fraction of cytoplasmic area (excluding nuclei) occupied by indicated glial cells in the GCL and ACL from WT and NP animals. Columns, mean and whiskers, standard deviation. Asterisks indicate statistically significant change due to NPC1 deficiency (**, p < 0.01; Wilcoxon signed rank test; WT: n = 6 mice; NP n = 5).

### Reduction of cholesterol accumulation in retinal neurons following direct delivery of CD

Next, we used intravitreal injections to study how CD impacts cholesterol accumulation in retinal neurons most affected by NPC1 deficiency *in vivo*. As control, contralateral eyes were injected with vehicle (PBS). Filipin staining of retinal sections showed that within 24 and 48 hours a single injection of CD reduced the extent of cholesterol accumulation in neurons of the ACL and GCL from mutant mice compared to vehicle (Fig. 3A, B). Inspection of NPC1-deficient neurons by TEM revealed that CD reduced the density of lamellar inclusions and induced the presence of inclusions bearing few lamellae or lacking them altogether (Fig. 3C). The quantitative analysis indicated layer-specific neuronal reactions to CD: in the GCL full inclusions decreased and half-full and empty counterparts appeared, whereas in the ACL mostly half-full inclusions were visible at both time points studied. Notably, ultrastructural features observed in the GCL indicated the release of inclusions from neurons *in vivo* (Fig. 3E) thus supporting our hypothesis based on *in vitro* data (Demais et al., 2016).

**Fig. 3.**
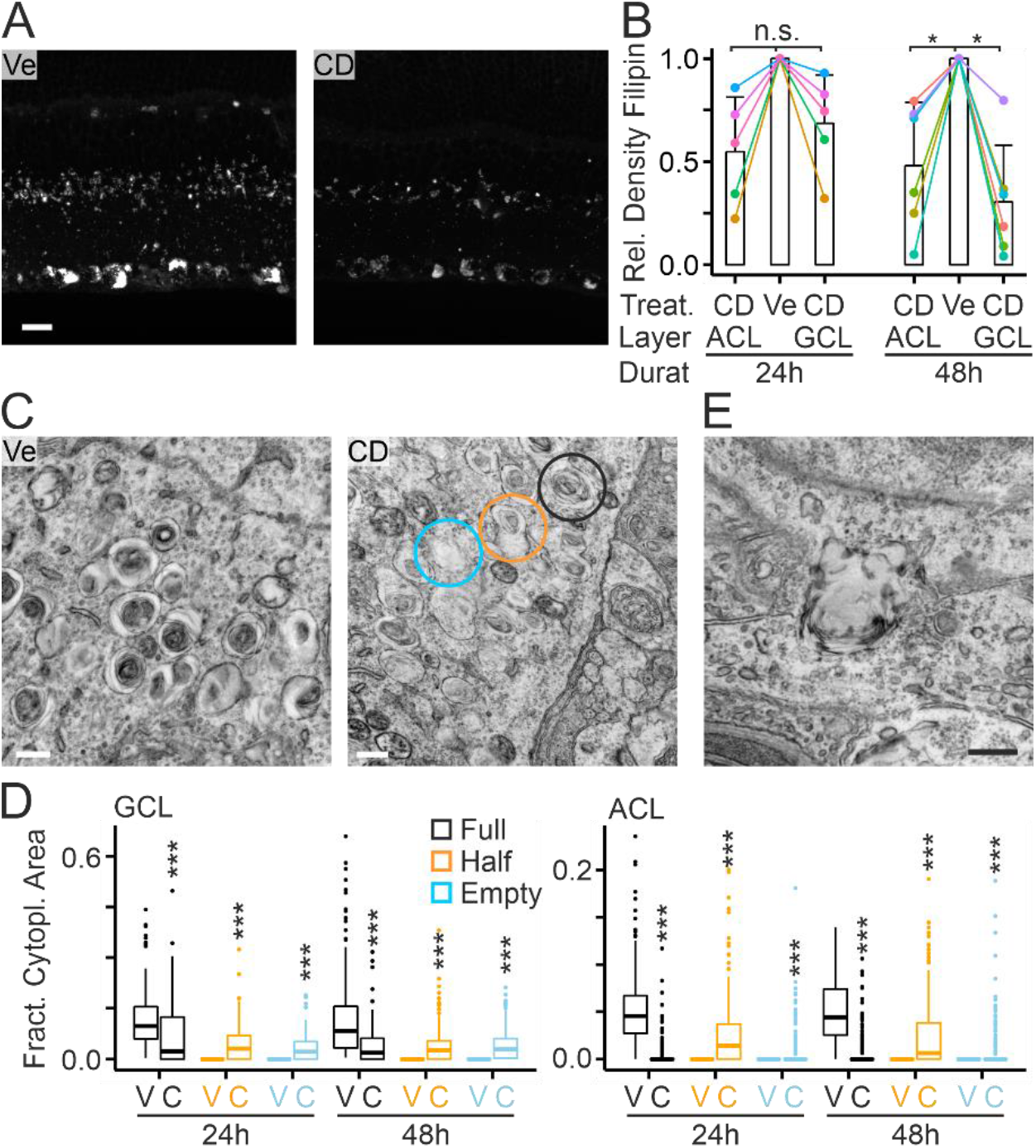
CD-induced reduction of cholesterol accumulation in NPC1-deficient retinal neurons. A, Fluorescence micrographs of filipin-stained retinae from NPC1-deficient mice 24h after intravitreal injections of PBS (Ve, vehicle) or CD. Scale bar: 25 μm. B, Effect of CD on the density of filipin-positive puncta in neuronal somata of the indicated layers from individual NPC1-deficient mice normalized to vehicle-treated contralateral eyes. Values were measured at indicated time periods after injections. Asterisks indicate statistically significant changes (p < 0.05; Wilcoxon signed rank test). C, Representative electron micrographs of lamellar inclusions found in RGCs of NPC1-deficient mice at 24h after intravitreal injection of vehicle and CD. CD treatment induced the occurrence of inclusions without lamellae (empty, sky blue) and with lamellae filling half of their interior (half, orange) in NP mice. Lamellar inclusions were absent from WT retinae. Inclusions with lamellae filling the entire interior are also indicated (full, black). Scale bar: 500 nm. D, Boxplots showing the fractional cytoplasmic area (bottom) of the different types of inclusions in neuronal somata of the GCL [left, 24h Vehicle 4 animals / 192 cells; CD: 4 / 124; 48h: V: 6 / 261; C: 5 / 171] and the ACL (right, 24h: V: 4 / 311; C: 4 / 206; 48h: V: 6 / 327; C: 5 / 284). CD-induced changes in the fractional cytoplasmic area covered by different types of inclusions were statistically significant compared to vehicle-treated eyes (***, p < 0.001; Kruskal-Wallis test with Dunn posthoc BH adjusted). E, Electron micrograph indicating the release of a laminar inclusion from a RGC after intravitreal injection of CD. Scale bar: 500 nm.

We hypothesized that CD removes cholesterol from inclusions thus permitting its partial incorporation into the cellular pool. Such action would require that CD reaches the interior of lamellar inclusions. To locate CD in subcellular compartments, we coupled CD to 2.5 nm gold nanoparticles. These particles can be visualized by TEM, and they have been used extensively in eye research (Masse et al., 2019). Elementary analysis of CD-AU particles by energy-dispersive X-ray spectroscopy confirmed the presence of carbon and gold (Fig. 4A). High resolution TEM showed the presence of monodispersed particles with a narrow size distribution (radius 2.6 ± 0.7 nm; n= 1000; Fig. 4B). *In vitro* tests on cultured RGCs confirmed that CD-AU particles reversed U18666A-induced accumulation of cholesterol as efficiently as CD alone (Fig. 4C). TEM inspection of retinae revealed that CD-AU particles were present in lamellar inclusions of RGCs already starting from 2h after intravitreal injection in NPC1-deficient mice suggesting their uptake via the endosomal-lysosomal system (Fig. 4D). We also found particles in the cytoplasm of Müller cells and in blood vessels at 24h after injections (Fig. 4E) in NP mice.

**Fig. 4.**
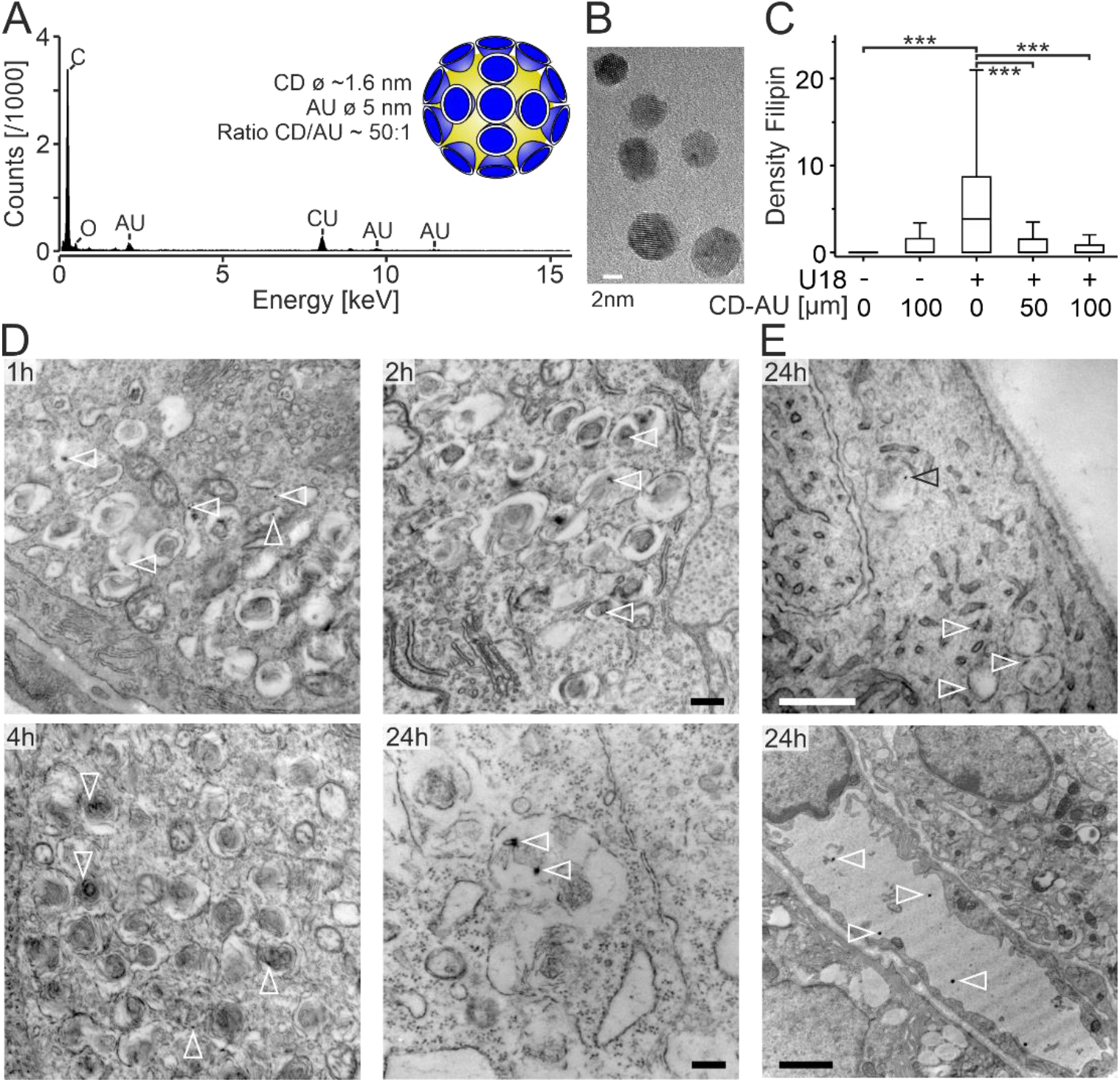
Tracking the subcellular localization of CD with gold. A, Elementary composition of CD-AU particles as shown by energy-dispersive X-ray spectroscopy. Inset, schematic view of CD-AU particles (approximately to scale). B, Electron micrograph of CD-AU particles. C, Bloxplots showing the density of filipin-positive puncta in somata of highly purified RGCs cultured under indicated conditions (-U18/CD-AU 0: n = 191 cells; -U18/CD-AU 100: 129; +U18/CD-AU 0: 215; +U18/CD-AU 50: 194; +U18/CD-AU 100: 178). Asterisks indicate statistically significant changes (***, p = 0; Kruskal-Wallis test with Dunn posthoc BH adjusted). Concentrations of CD-AU indicated in μM. Cholesterol accumulation was induced by U18666A. D and E, Electron micrographs showing the presence of CD-AU (white arrowheads) in lamellar inclusions of RGCs (D), in the cytoplasma of Müller cells (E, top) and in the lumen of blood vessels (E, bottom) at indicated times following intravitreal injection in NPC1-deficient mice. Scale bars: 500 nm (D; E, top), 2 μm (E, bottom).

### Reaction of retinal glia to intravitreal administration of CD

Next, we studied how glial cells react to intravitreal injections of CD in WT and NP mice. Immunohistochemical staining of retinal whole-mounts revealed an increased density of GFAP- and IBA-positive cells 48h after injection of CD, but not of vehicle. Notably, these changes were only observed in NP, but not in WT animals (Fig. 5A, B). TEM inspection showed layer- and cell type-specific responses of glial elements to CD injections in NP mice. CD strongly increased the fractional area covered by glial cells with clear cytoplasm in the GCL and ACL of NP mice at 24 and 48h after injections (Fig. 5C, D). Glial cells with dark cytoplasm reacted similarly, but the expansion was more marked at 48h after the injection (Fig. 5C, D). Notably, Müller cells from NP mice did not show any changes in fractional area in response to injections (Fig. 5C, D). Moreover, no changes in area covered by glial elements were observed in WT mice or after vehicle injections.

**Fig. 5.**
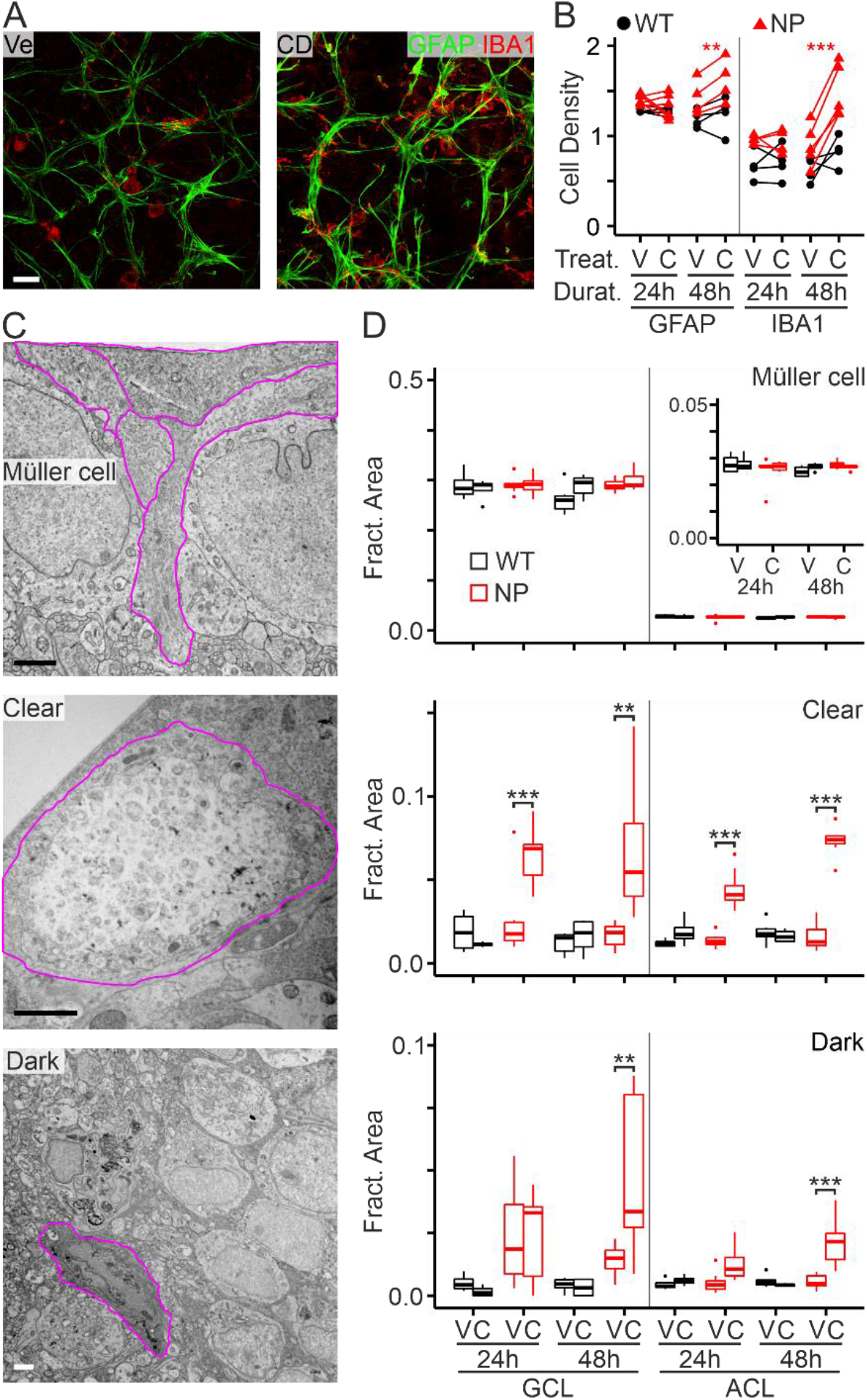
Reaction of retinal glial cells to intravitreal injections. A, False-color fluorescence micrographs of retinal whole-mounts from NPC1-deficient mice 48h after intravitreal injection of PBS (Ve, vehicle) or CD and immunohistochemical staining for Gfap- (green) and Iba- (red) positive cells. Scale bar: 25 μm. B, Density of Gfap- and Iba-positive cells in retinal whole-mounts of WT and NP mice at indicated times after intravitreal injection of vehicle (V, left eye) or CD (right eye). CD induced statistically significant changes after 48h in NPC1-deficient mice (**, p<0.01; ***, p < 0.001; paired t-test). C, Representative electron micrographs of Müller cells and non-neuronal cells with clear and dark cytoplasm (outlined by magenta lines) in retinae of NP mice after intravitreal injection of CD. Scale bar: 2 μm. D, Boxplots of area covered by indicated glial cells as fraction of total area analysed in the retinal GCL and ACL per mouse with indicated genotypes at indicated times after intravitreal injections of vehicle (V, left eye) or CD (C, right eye) (24h WT Vehicle / CD: n = 4 / 4 animals, NP V / C: 10 / 9; 48h WT V / C: 6/6, NP V / C: 8 / 7). Inset, fractional area of Müller cells in ACL at modified scale. Asterisks indicate statistically significant changes induced by CD (**, p < 0.01; ***, p < 0.001; ANOVA with Tukey’s post hoc test for each treatment duration).

### CD-induced presence of lamellar inclusions in glial cells

Our previous work *in vitro* suggested that CD enables neurons to release inclusions into the extracellular space (Demais et al., 2016). Sporadically, we observed ultrastructural features in the GCL that indicated the occurrence of release *in vivo* (Fig. 3E), but the fate of these structures remained unclear. The lamellar inclusions released from neurons may have been taken up by glial cells. Histochemical and immunohistochemical staining of unesterified cholesterin and glial cells, respectively revealed a colocalization of filipin-stained cholesterol in GFAP- and IBA1-positive cells in retinae from NP-deficient mice following intravitreal injections of CD, but not of vehicle (Fig. 6A). Inspection of retinae by TEM revealed that CD strongly increased the size of phagosome-like structures in both clear and dark glial cells in the GCL and ACL of NP animals and that these phagosomes contained exclusively lamellar inclusions (Fig. 6B, C). These changes were not induced by vehicle and they were absent from WT animals (Fig. 6C). Together, our results indicated that CD induces an uptake of lamellar inclusions by glial cells leading to a strong expansion of the area covered by glial elements.

**Fig. 6.**
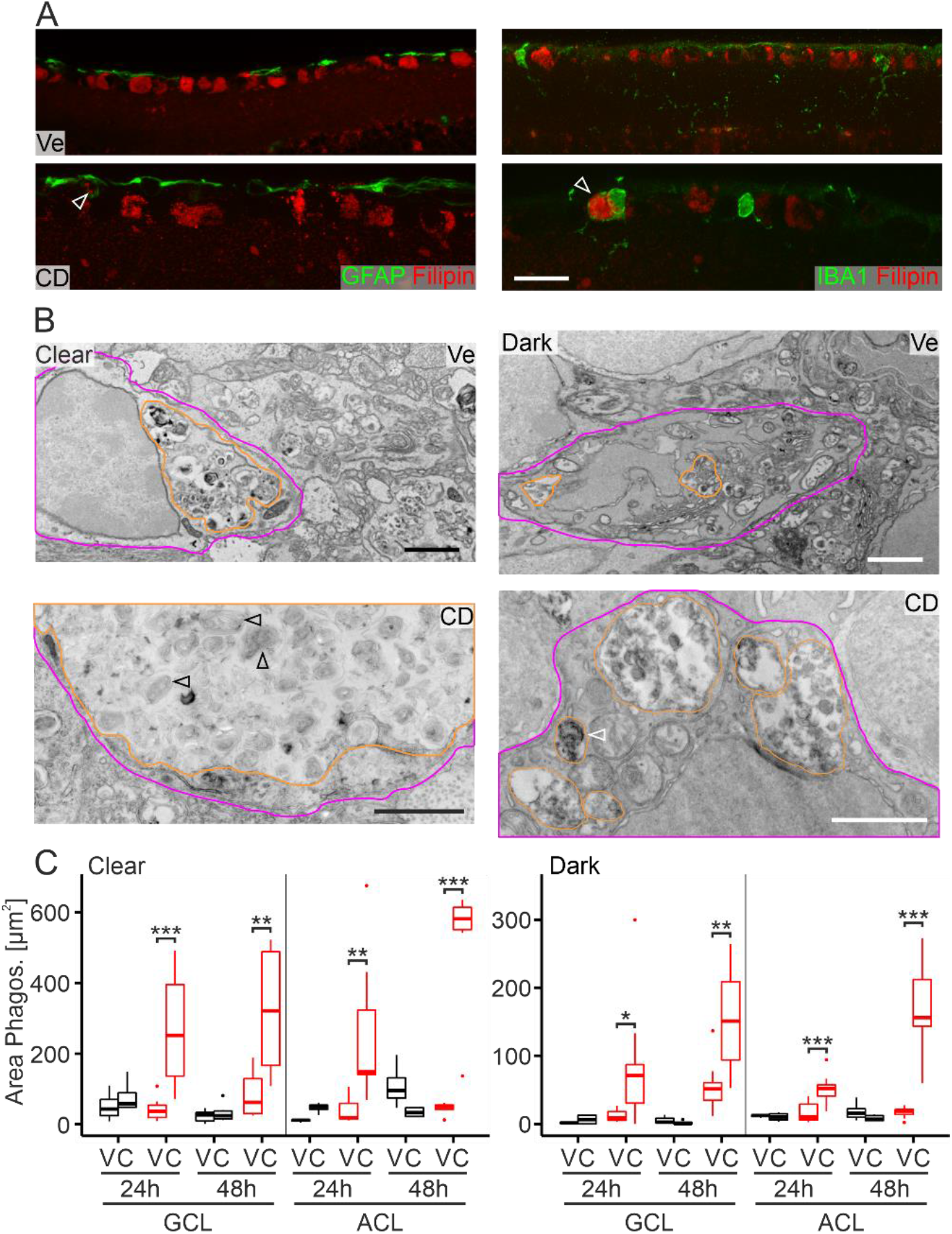
CD-induced uptake of lamellar inclusions by glial cells. A, Representative false-color fluorescence micrographs showing the presence of filipin-stained unesterified cholesterol in somata of GFAP- (left) and IBA-positive (right) cells in the GCL of retinae from mice at 24h after intravitreal injections of CD. No filipin staining was observed in retinal glial cells from vehicle-injected contralateral eyes. Scale bar: 25 μm. B, Representative electron micrographs of phagosome-like structures (orange outline) in clear (left; magenta outline) and dark (right; magenta outline) glial cells in retinae from NP mice at 48h after intravitreal injection of vehicle (top) and CD (bottom). Note that following CD injections, phagosomes were strongly enlarged and contained almost exclusively lamellar inclusions. Scale: 500 nm. C, Boxplots showing the area covered by phagosomes in clear (left) and dark (right) glial cells in the retinal GCL and ACL from WT and NP animals at indicated times after intravitreal injections of vehicle (V, left eye) or CD (C, right eye) (24h WT Vehicle / CD: n = 4 / 4 animals, NP V / C: 10 / 9; 48h WT V / C: 6 / 6, NP V / C: 8 / 7). Asterisks indicate statistically significant changes induced by CD compared to vehicle controls at each treatment duration (*, p > 0.5; **, p < 0.01; ***, p < 0.001; ANOVA with Tukey’s post hoc test).

### CD-induced presence of neutrophil granulocytes

Observation of retinal sections from CD-injected NP mice by TEM revealed the presence of cells with characteristic features of neutrophil granulocytes. These features include a diameter of 12-15 μm, polymorph nuclei rich in heterochromatin and granule-rich cytoplasm (Duvvuri et al., 2020; Yipp et al., 2012) (Fig. 7). To follow up on this, we performed immunohistochemical stainings of retinae with MPO, a marker of neutrophil cells, and CD68 to discern microglial cells (Fig. 7B). These experiments showed that CD strongly enhanced the density of MPO-positive and CD68-negative cells in NP mice, whereas no changes were observed in vehicle-injected animals or in WT mice (Fig. 7C). Notably, neutrophils seemed predominantly located in the nerve fiber layer and GCL. Quantitative ultrastructural analysis confirmed the appearance of neutrophils and their layer-specific localization upon CD injection (Fig. 7D). These findings suggested that CD induces the entry of neutrophils into retinae of NP mice.

**Fig. 7.**
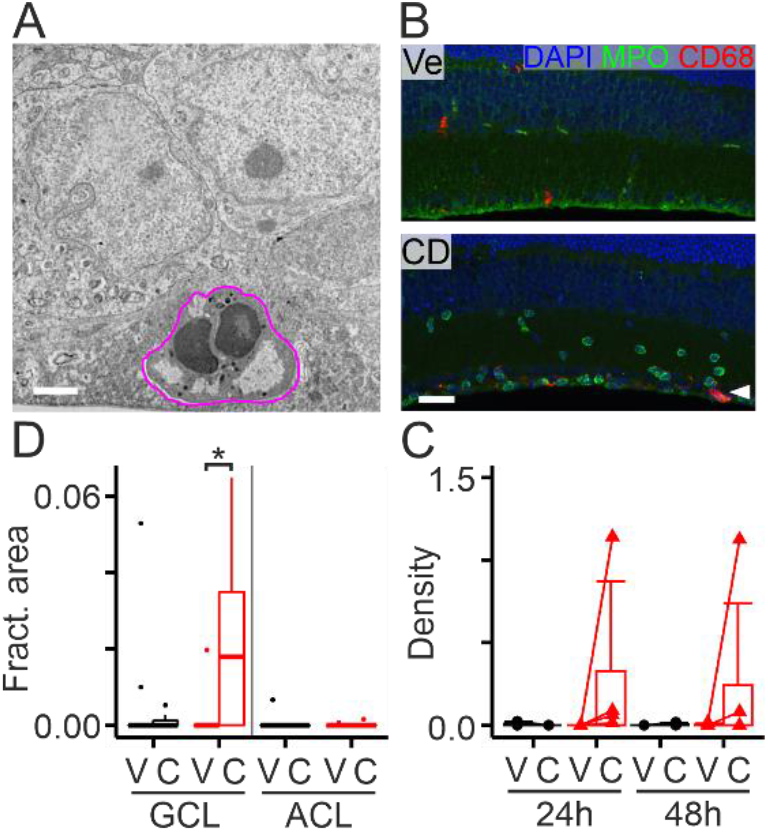
CD-induced presence of neutrophils in retinae of NPC1-deficient mice. A, Electron micrograph of a cell in the nerve fiber layer (outlined in magenta) with typical ultrastructural features of neutrophils. Retinal section from NP mouse 48h after intravitreal injection of CD. Scale bar: 2 μm. B, False-color fluorescence micrographs showing the distribution of MPO-positive cells (green), CD68-positive microglia (red) and of DAPI-positive nuclei (blue) in retinal sections from NP mice after intravitreal injection of vehicle (Ve) and CD. Scale bar: 50 μm. Arrowhead indicates GCL. C, Plots showing individual and average density of MPO-positive/CD68-negative cells in retinal sections from WT and NP mice at indicated times after intravitreal injections of vehicle (V) or CD. The differences between CD versus vehicle were not statistically significant (24h WT: n = 3, p = 0.422; NP: n = 4, p = 0.344; 48h WT: n = 4, p = 0.391; NP: n = 5, p = 0.331; paired t test). D, Boxplots showing the fractional area occupied by neutrophil-like cells in the retinal GCL and ACL from WT and NP animals at 24 or 48h after intravitreal injections of vehicle (V, left eye) or CD (C, right eye) (WT Vehicle / CD: n = 10 / 10 animals, NP V / C: 17 / 16). The asterisk indicates a statistically significant change induced by CD (*, p < 0.05; ANOVA with Tukey’s post hoc test).

## Discussion

Using the mouse retina as experimental model and intravitreal injections as mode of drug and vehicle administration we provide new insight how CD reverts cholesterol accumulation in NPC1-deficient neurons *in vivo*. Our results indicate that CD is rapidly taken up by neurons and enters lamellar inclusions. This was shown by new CD-AU conjugates enabling ultrastructural detection of CD. The conjugates are likely to behave similarly to CD. Previous reports showed cellular uptake of cyclodextrins coupled to dextran (Rosenbaum et al., 2010), a bodipy fluophore (Dai et al., 2017), fluorescein (Plazzo et al., 2012) and fluorescent nanoparticles (Donida et al., 2020) *in vitro*. Our control experiments confirmed similar efficacy of CD-AU and CD in reducing the cholesterol accumulation in NPC1-deficient neurons. CD reduced the density of lamellae inside inclusions suggesting that the compound enables a partial exit of cholesterol and possibly other lipids from these structures independently from NPC1 (Abi-Mosleh et al., 2009). This may involve alternative shuttle mechanisms (Heybrock et al., 2019). Our observation that CD acts inside cells is supported by findings *in vitro* that the molecule reduces cholesterol accumulation with cell-specific delays (Dai et al., 2017; Meske et al., 2014) and that endocytotic uptake is required for its action (Rosenbaum et al., 2010; Vance and Karten, 2014). The continued presence of lamellae inside inclusions may be caused by *de novo* formation or by lipids that are not affected by CD. The layer-specific differences in the size and density of lamellar inclusions and in the reaction to CD treatment suggest neuron-specific modes or rates of lipid accumulation and of their CD-mediated clearance.

Previous studies on NPC1 mutant cells proposed that cyclodextrin induces the secretion of cholesterol *in vitro* (Chen et al., 2010; Feltes et al., 2020; Ilnytska et al., 2021; Juhl et al., 2021; Sedgwick et al., 2018; Vacca et al., 2019) probably by distinct modes depending on cell type and experimental conditions. Our observations on primary cultures suggested that CD enables neurons to release the bulk of lipid-laden lamellar inclusions into the extracellular space (Demais et al., 2016). Our new findings *in vivo* support this hypothesis. The appearance of extravasate neutrophils and the presence of lamellar inclusions in retinal glial cells upon CD injection in mutant mice suggest that these structures were released by neurons and thereby triggered responses from non-neuronal cells. Why should neurons secrete these structures? This mode is probably the only way for neurons to prevent lipid overload. Defects in NPC1 or NPC2 protein enhance the intracellular pool of cholesterol in patient fibroblasts by ten-fold (Lange et al., 1998), and a similarly sized lipid burden can be expected in NPC1-deficient neurons. Neurons are probably unable to handle this large amount of lipids as they cannot esterify and store cholesterol (Aqul et al., 2011; Nieweg et al., 2009; Peake and Vance, 2012; Saadane et al., 2019; Zheng et al., 2012). Moreover, their unique pathway to eliminate surplus cholesterol by CYP46A1 may become saturated (Lund et al., 1999; Lutjohann et al., 1996; Pfrieger and Ungerer, 2011). In the retina, this enzyme is expressed by RGCs (Bretillon et al., 2007; Nieweg et al., 2009; Pikuleva and Curcio, 2014; Zheng et al., 2012). These assumptions are supported by findings that intra-cerebroventricular injection of CD in NPC1-deficient mice did not affect cholesteryl ester synthesis or Cyp46a1 expression in the brain (Aqul et al., 2011). The CD-induced enhancement of 24S-hydroxycholesterol levels in the cerebrospinal fluid of NPC patients was probably due to decreased loss of neurons (Ory et al., 2017).

The observation of strongly enlarged phagosomes filled with lamellar inclusions in glial cells suggests that these cells handle the superfluous material released from neurons following CD administration. Several studies reported that CD attentuates glial activation (Aqul et al., 2011; Cougnoux et al., 2018; Liu et al., 2009; Maulik et al., 2012; Meyer et al., 2018; Park et al., 2019; Ramirez et al., 2010), but these observations were made weeks after drug injections and thus reflect the steady-state situation. We suggest that astrocytes and microglial cells bearing clear and dark cytoplasm take up inclusions, respectively. These cells are known to clear debris and surplus lipids from the healthy and diseased brain (Chausse et al., 2020; Galloway et al., 2019; Loving and Bruce, 2020; Márquez-Ropero et al., 2020; Prinz et al., 2019; Tremblay et al., 2019; Yang et al., 2021). Recent studies suggested that phagocytotic activity of microglial cells is affected by NPC1 deficiency *in vivo* (Boyle et al., 2020; Colombo et al., 2021; Kavetsky et al., 2019) and *in vitro* (Peake et al., 2011; Stein et al., 2012), but these changes are probably overcome by CD (Colombo et al., 2021; Peake and Vance, 2012). Notably, Müller cells, which can be readily identified by their location and ultrastructural features, seemed largely unaffected by NPC1-deficiency at the presymptomatic age and they showed no discernable ultrastructural changes in response to CD injection. This observation underlines the exquisite specialisation of glial cells in the retina (Vecino et al., 2016).

Taken together, our data suggest that the CD-induced reversal of cholesterol accumulation in NPC1-deficient retinal neurons is a non-cell autonomous process requiring the coordinated interactions of different types of glial cells.

## Author contributions

AB: Data Curation, Methodology, Investigation, Validation, Visualization, Writing - Review & Editing. VD: Data Curation, Methodology, Investigation, Validation, Visualization, Writing - Review & Editing. ICS: Methodology, Investigation, Resources, Validation, Visualization, Writing - Review & Editing. TH: Methodology, Investigation,Writing - Review & Editing. MR: Methodology, Resources, Writing - Review & Editing. VP: Methodology, Investigation, Visualization, Writing - Review & Editing. MP: Methodology, Investigation. SR: Project administration, Resources, Writing - Review & Editing. FWP: Conceptualization, Data Curation, Formal analysis, Funding acquisition, Project administration, Resources, Software, Supervision, Validation, Visualization, Writing - original draft, Writing - Review & Editing.

## Acknowledgments

We thank Yvonnick Pongérard and Nolwenn Couqueberg for help with the management of the mouse colony.

